# Emergence of complex dynamics of choice due to repeated exposures to extinction learning

**DOI:** 10.1101/2020.04.17.046136

**Authors:** José R. Donoso, Julian Packheiser, Roland Pusch, Zhiyin Lederer, Thomas Walther, Metin Uengoer, Harald Lachnit, Onur Güntürkün, Sen Cheng

## Abstract

Extinction learning, the process of ceasing an acquired behavior in response to altered reinforcement contingencies, is not only essential for survival in a changing environment, but also plays a fundamental role in the treatment of pathological behaviors. During therapy and other forms of training involving extinction, subjects are typically exposed to several sessions with a similar structure. The effects of this repeated exposure are not well understood. Here, we studied the behavior of pigeons across several sessions of a discrimination learning task in context A, extinction in context B, and a return to context A to test the context-dependent return of the learned responses (ABA renewal). By focusing on individual learning curves across animals, we uncovered a session-dependent variability of behavior: (1) During extinction, pigeons preferred the unrewarded alternative choice in one-third of the sessions, predominantly during the first one. (2) In later sessions, abrupt transitions of behavior at the onset of context B emerged, and (3) the renewal effect decayed as sessions progressed. We show that the observed results can be parsimoniously accounted for by a computational model based only on associative learning between stimuli and actions. Our work thus demonstrates the critical importance of studying the trial-by-trial dynamics of learning in individual sessions, and the power of “simple” associative learning processes.

## Introduction

Animals modify their behavioral repertoire based on the consequences of their own actions. Whether a certain behavior is reinforced or not, respectively increases or decreases the likelihood that this behavior is repeated in a similar situation (Skinner 1938). This process of operant conditioning is not only pivotal for the discovery of purposeful actions, but also plays a fundamental role in the development of pathological behaviors such as drug addiction, overeating and gambling (Bouton 2011; Everitt and Robbins 2005; Hyman 2005; Kelley 2004; Redish 2004). These pathological behaviors might arise from reinforcement for operant actions such as approach, handling and consumption, which are under the control of natural stimuli such as, for example food, drug, games, and money, and context such as, for instance a restaurant, drug den or casino. Once a behavior is acquired, the ability to extinguish it as a result of altered reinforcement contingencies is also essential for survival. This so-called extinction learning is defined as the reduction of conditioned responses to a previously reinforced stimulus when that stimulus no longer signals reinforcement. The importance of extinction learning is emphasized by the fact that all vertebrate and invertebrate species tested exhibit this ability (Barad et al. 2006; Eisenberg and Dudai 2004; Galatzer-Levy et al. 2013; Gao et al. 2018; Gottfried and Dolan 2004; Lengersdorf et al. 2014; Milad et al. 2007; Stollhoff et al. 2005). Extinction learning is also relevant during therapy and other learning settings, where, for example, patients are trained to withdraw undesired behaviors, or when children are required to suppress recurrent spelling mistakes.

Although extinction may involve some erasure of the previously acquired memory (McClelland and Rumelhart 1985; Rescorla and Wagner 1972), there is strong evidence that it also involves new learning (Bouton 2019; Todd 2013; Trask et al. 2017). According to the latter view, during extinction, the previously acquired memory trace responsible for a particular behavior is inhibited by a secondary memory trace. This new learning seems to depend on the context for expression, which is compatible with the idea that contextual cues can support memory retrieval (Tulving and Thomson 1973). A prominent phenomenon in support of extinction as new learning is the renewal effect, where an extinguished behavior reemerges when subjects are removed from the context where extinction learning took place (Bernal-Gamboa et al. 2017; Bouton 2019; Nieto et al. 2017, 2020; Todd 2013; Todd et al. 2014). This context-dependence of extinction has a severe downside under conditions like exposure therapy, where a seemingly extinguished pathological behavior resurfaces when patients switch from a therapy context to their regular environment (Conklin and Tiffany 2002). Therefore, understanding extinction and the factors that influence the reappearance of the extinguished behavior can be helpful in developing treatments for several pathological behaviors and in preventing their reappearance (Bouton 2011; Conklin and Tiffany 2002).

Another relevant aspect of extinction learning is that it is not limited to a decrease in previously reinforced responses, but it can also drive the emergence of new, previously non-reinforced behaviors (Antonitis 1951; Fuller 1949; Grow et al. 2008; Lattal and Lattal 2012; Tinsley et al. 2002). This side-effect of extinction has been exploited to shape behavior. For example, when the extinction of problematic behaviors drives the emergence of socially acceptable responses, the latter can be reinforced in order to replace undesired behaviors with desired ones (Grow et al. 2008). In the same way, extinction-induced variability becomes particularly relevant in real world settings and experimental designs where there are multiple alternative choices in a particular situation. While there are studies of extinction in the presence of alternative responses (e.g., Winterbauer and Bouton 2010), research has focused mostly on the cessation of target responses, often neglecting the effect that extinction might have on other available actions (André et al. 2015; Méndez-Couz et al. 2019).

Experiments involving extinction learning, and/or testing for the renewal effect, rarely involve more than one conditioning-extinction sequence (e.g., Bernal-Gamboa et al. 2017; Bouton 2019; Nieto et al. 2017, 2020; Todd et al. 2014; Trask et al. 2017); but see (Anger and Anger 1976; Bullock 1960; Jenkins 1961). Outside the laboratory, however, an animal’s learning history comprises multiple instances in which a period of conditioning is followed by extinction. Such multiple conditioning-extinction sequences could differ in the stimuli involved, but may share a similar structure. Nevertheless, in most studies involving repeated extinction, the same association that is learned and extinguished in a given session is relearned and re-extinguished in subsequent sessions (Bai and Podlesnik 2017; Craig and Shahan 2016; Craig et al. 2019; Podlesnik and Sanabria 2011). In spite of its relevance, the effect of this repeated exposure to the same task structure with varying learning content has not been studied systematically. Another aspect that remains to be elucidated is the effect of the learning history both on the emergence of alternative behaviors, and on the prevalence of the renewal effect across sessions.

In the present study, we focus on the variability of choice behavior, which is a specific aspect of the more general behavioral variability that animals can express under altered reinforcement contingencies (Eckerman and Lanson, 1969; Neuringer et. al., 2001). Specifically, we analyzed the choice behavior of pigeons undergoing multiple sessions of a discrimination learning task (Fig. 1). Each session consisted of three subsequent stages: (1) learning of correct responses to two novel visual stimuli in acquisition context A; (2) extinction learning for one of the learned stimulus-response mappings in context B, and (3) return to context A to test for ABA renewal of the extinguished mapping (e.g., Nieto et al. 2017). We used pigeons due to their ability to withstand multiple sessions of learning of more than 1000 trials per day. Moreover, their slow pace of learning allowed us to focus on the trial-by-trial dynamics of choice behavior in single sessions and individual subjects. This approach uncovered a rich repertoire of choice behavior associated with the extinction and renewal-test phases that are key to understanding associative learning mechanisms. Additionally, we provide a parsimonious model consisting of an associative network and a winner-takes-all decision-making process, where the action unit with the highest activation drives behavior. This simple model could account for several aspects of the behavioral phenomena observed in the data.

**Figure 1.**
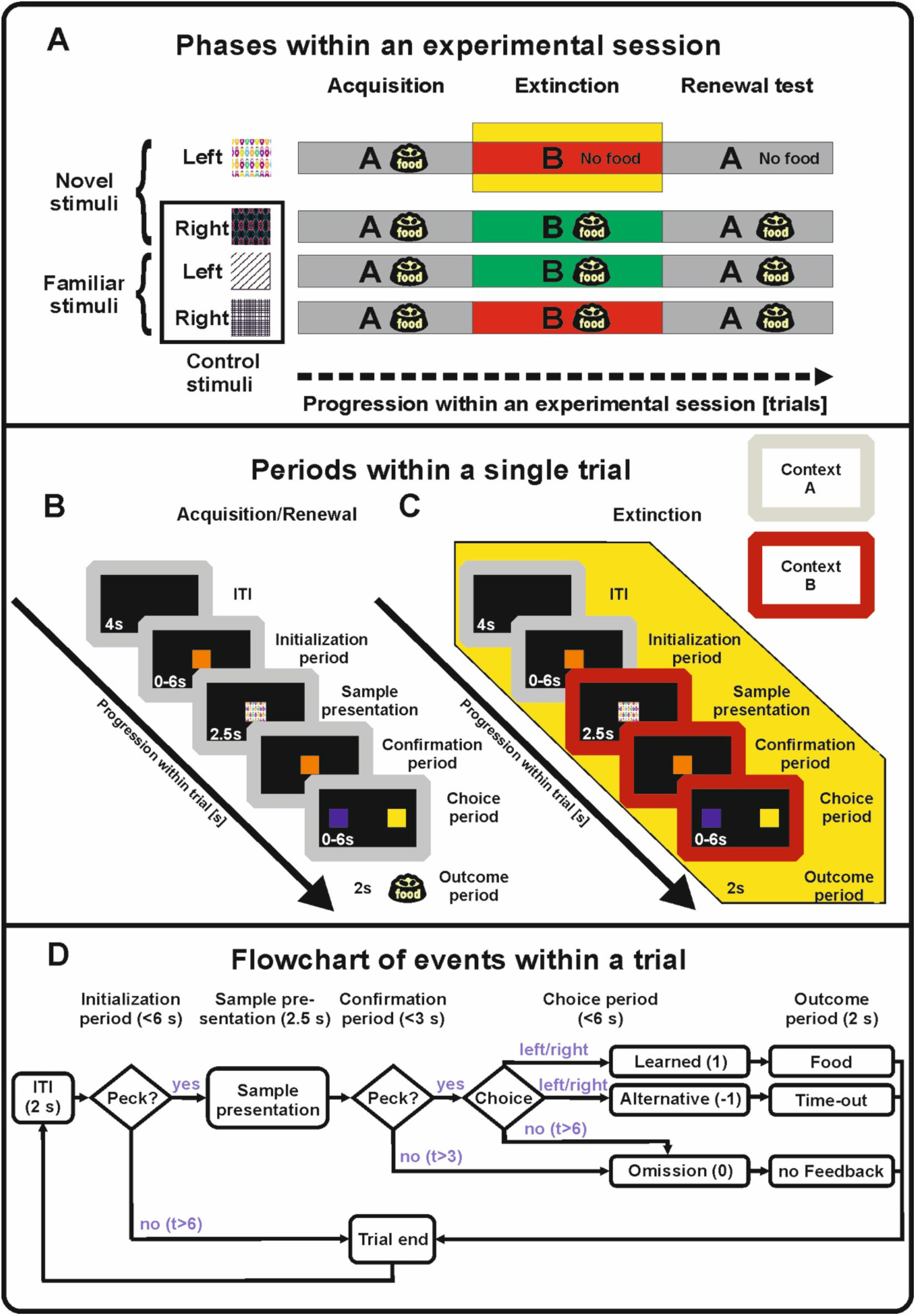
Experimental design. (A) Overview of the behavioral paradigm. During the *acquisition phase* (context A) animals learned to associate left/right choices with two novel stimuli. After reaching a learning criterion, extinction started in context B. One novel stimulus was randomly assigned to be extinguished (highlighted in yellow). In the *renewal test phase* context A was re-established, while pecking the extinction stimulus still did not yield reward. (B) Trial structure and sequence of screen presentations during acquisition/renewal test. The transitions between the trial periods are described in panel D. (C) Trial structure and sequence of screen presentations during extinction trials. Initialization peck on the center screen triggered a change in the light conditions to context B (red frame) 1 second before the stimulus presentation, which remained until the end of the trial. In extinction stimulus trials the outcome period remained void of any feedback regardless of the animal’s decision (highlighted in yellow). (D) Flowchart depicting transitions between trial periods and possible outcomes. Note that in trials involving the extinction stimulus trials during the *extinction phase* and the *renewal-test phase*, the outcome period remained void of any feedback regardless of the animal’s decision (cf. panel C).

## Materials and Methods

The behavioral data reported here were obtained in the same experiment in which electrophysiological data were collected to study the neural correlates of the reward prediction error in extinction learning and renewal (Packheiser et al. 2020a). The present study is independent from the previous study regarding focus, hypotheses, data analysis and interpretation. Specifically, the previous study did not analyze learning curves in individual sessions or across repeated extinction sessions. Data collection for the present study started after surgical procedures were completed.

### Subjects

Eight experimentally naive, adult pigeons (*Columba livia*) of unknown sex and age obtained from private breeders were used as subjects in the present experiment. Birds were housed in individual wire-mesh cages or local aviaries within a colony room. The housing facilities were controlled for light cycles (12 h light/dark cycles starting at 8 am), temperature and humidity. All animals had *ad libitum* access to water and were kept between 80% and 90% of the free-feeding body weight. The food deprivation was necessary to keep the animals engaged in the experimental procedures. All animals were treated in accordance with the German guidelines for the care and use of animals in science. The experimental procedures were approved by a national ethics committee of the State of North Rhine-Westphalia, Germany and were in agreement with the European Communities Council Directive 86/609/EEC concerning the care and use of animals for experimental purposes. Pigeons were part of an electrophysiological experiment in which single-units were recorded over multiple sessions of learning. Prior to session onset, we checked whether neural signals were present and could be isolated on the channels. If so, the animals were tested and a behavioral session was generated. In the results section, pigeons were enumerated according to the number of sessions they contributed: 1-3 (26 sessions), 4 (23 sessions), 5 (22 sessions), 6 (13 sessions), 7 (11 sessions), 8 (6 sessions).

### Apparatus

The experimental procedures were conducted in custom-made Skinner boxes (35cm x 35cm x 35cm situated in sound-attenuating cubicles (80cm x 80cm x 80cm)(Packheiser et al. 2020b). Each Skinner box featured three rectangular pecking areas (5cm x 5cm) that were horizontally arranged on the rear wall. Depending on the type of Skinner box, either touch screens or translucent response keys combined with a mounted LCD flat screen monitor were used to track pecking responses. A feeder was located below the central pecking site (feeding area: ~1cm x 1cm) to deliver food rewards during the experiments. White LED strips mounted to the ceiling were used to illuminate the experimental chamber. Furthermore, red and green LED strips were attached to the ceiling to enable flexible contextual changes during the paradigm. If the animals successfully pecked onto a response key, an auditory feedback sound was presented. The hardware was controlled by a custom written MATLAB program (The Mathworks, Natick, MA, USA) using the Biopsychology toolbox (Rose et al. 2008).

### Procedure

Pigeons underwent 6 to 26 sessions of a discrimination learning task in which animals were subject to acquisition, extinction and a renewal-test phase within one session (Fig. 1A) (Packheiser et al. 2019; Starosta et. al. 2014). In each trial, animals were presented one visual stimulus. In response, the animal had to make either a left or a right choice at the end of the trial depending on the stimulus identity. During a session, the animals were confronted with four different stimuli presented in a pseudorandomized order. Two of the stimuli were associated with a left choice and the other two stimuli were associated with a right choice. Animals were pre-trained on two of the stimuli prior to the experimental sessions studied here. The precise nature of the pre-training procedure is explained in detail in (Packheiser et al. 2020a). Hence, two of these stimuli were familiar to the animals and served as control stimuli as well as fix points during the experiment. The other two stimuli were session-unique and the stimulus-response mapping had to be learned in the acquisition phase through trial-and-error. For the familiar stimuli, we used black and white stimuli featuring either a squared or a striped texture (see Figure 1A). Novel stimuli were always colored and featured different forms or shapes to allow for a discrimination both in color and spatial features.

#### Acquisition

Figure 1B shows a sequence of screen presentations during acquisition. In brief, each trial started with the presentation of an initialization key for up to 6s. A successfully registered key peck to the center response key triggered the sample presentation. One of four stimuli (see below) was presented for 2.5s on the center key. Following the stimulus presentation, the animals were required to confirm that they attended the target stimulus by pecking on the center key once more. After pecking on the confirmation key, the center key stimulus disappeared and the two choice keys were illuminated (a blue one on the left and a yellow one on the right, counterbalanced across animals). The animal had to decide on a left or a right choice depending on the identity of the stimulus that was presented earlier. If the animals made the correct choice (henceforth ‘learned choice’), a 2s long reward period commenced during which the food hopper was illuminated, a sound occurred and food was available. In the case of an incorrect choice (henceforth ‘alternative choice’), a different sound occurred and the lights in the chamber were turned off for 2s as a mild punishment. Consecutive trials were separated by an inter-trial-interval (ITI) of 4s duration. A detailed description of the events and transitions between periods within a single trial is depicted as a flowchart in Figure 1D. The acquisition phase comprised a minimum of 150 trials and ended once the animals satisfied all of the following criteria: the animals initialized 85% of the trials correctly, performed above 85% correctly in response to the novel stimuli and above 80% correctly is response to the two familiar stimuli. Performance values were calculated as a running average over the past 100 trials.

#### Extinction

The subsequent extinction phase was marked by two key differences as compared to the acquisition phase (Fig. 1C). (1) One of the novel stimuli was randomly chosen as the extinction stimulus, i.e., it was no longer followed by reward nor by punishment after any choice the animal made. Instead, the feedback phase was replaced by a 2s-long period void of feedback. (2) After the initialization of the trial by the animal, a colored LED light (indicator of context B) replaced the white house light used in the acquisition phase. The colored LED light was turned on one second before the stimulus presentation, and remained on until the end of the trial or until a punishment condition was met. We used two different light colors to signal the extinction context in order to ensure that the physical identity of the light was not driving behavioral effects in the extinction context. Within a given session during extinction training, both the red and green contexts were present. Each context-color was specifically associated with two experimental stimuli with contralateral response mappings, namely one familiar and one novel stimulus. For example, in a given session, when the novel-left peck mapping was extinguished under context B1, then a familiar-right peck mapping was still rewarded under the same context, and the two remaining stimulus-response mappings continue to be rewarded under context B2. Thus, in each session one of the contexts B1 or B2 was associated with extinction learning whereas the other context was not. Additionally, we also changed the audio cues for the correct and incorrect responses to add another modality change highlighting the contextual switch. The extinction phase comprised a minimum of 150 trials and ended when the following conditions were all met: the animals initialized 85% of the trials correctly, performed above 80% correctly in response to the novel non-extinction stimulus and more than 75% correctly in response to the two familiar control stimuli, and emitted the learned choice in response to the extinction stimulus less than 20% of the time. All performance values were calculated as a running average over the past 100 trials.

#### Renewal-test

Finally, the renewal-test phase was used to study the return of the learned choice when the context was switched back to the acquisition context A (Fig. 1A). Importantly, the extinction stimulus remained without feedback to measure the renewal effect. The renewal-test phase lasted for a fixed number of 250 trials and required no behavioral criterion to end. Its end also marked the end of the session. Data acquisition was ended when the animals no longer provided any neural data for the study by Packheiser et al. (2020a).

### Analysis

To quantify the preference for a target choice in a given phase, we counted the corresponding responses (*k*), and expressed them as a proportion of the total number of trials (*N*) in that phase (response rate = k/N). To assess the significance of a response count *k*, we calculated the probability *p*(*k*) of obtaining at least *k* responses by chance in *N* random trials under the null hypothesis. Since responses to the extinction stimulus are not rewarded during the extinction and renewal-test phases, our null-hypothesis assumes unbiased random responses, such that each one of the three possible outcomes (learned choice, alternative choice, or omission) can occur with probability ⅓. If the probability of observing at least *k* responses by chance was below a threshold of 0.05, we regarded the count as significant. This method, however, overlooks those cases where a non-significant number of responses are arranged in a chain of persistent responses, which is also unlikely to occur by chance. To consider those cases, we also measured the length *L* of the longest chain of persistent responses found on a specific phase (AP in Figs. 3C and 6A), and calculated the probability *p*(*L*) of obtaining a chain of at least *L* trials by chance in *N* random trials. Finally, the choice behavior was regarded as significant, if one of the aforementioned tests yielded a *p*-value below 0.05. We also used this measure to quantify the prevalence of a target response, which was defined as the proportion of animals expressing a significant number of target responses in a given session. To quantify the dependency of these response measures on the session number, we calculated the rank order correlation (Spearman’s r) between session number and both preference and prevalence. Since the number of animals contributing with data gradually decreased with session number, we grouped the data in blocks encompassing a comparable number of data points: Sessions 12 to 23 in blocks of two sessions (9 to 12 data points per block), and the data from sessions 24 to 26 in one single block (9 data points). The rank-order correlations (Spearman’s r) between the session number and the prevalence/prevalence of a particular response across animals was obtained from the grouped data unless stated otherwise. The p-values accompanying the reported correlations correspond to a t-test for the significance of the respective correlation coefficient (r).

To visualize the behavior in response to a specific stimulus within single sessions, we plotted the cumulative record of responses to that stimulus as a function of trial number (Fig. 3). For each trial, the choice behavior of the animal, namely, omissions, alternative choice, and learned choice, were encoded as 0, −1, and 1, respectively. To assess the presence of abrupt transitions of behavior upon the onset of the extinction phase, we focused on the responses to the first 5 presentations of the extinction stimulus under context B. In this case, our null hypothesis is that animals continue to emit the learned choice with at least 80% probability required to accomplish the acquisition phase (i.e., no behavioral change at the beginning of the extinction phase). Thus, if animals emit at least 3 non-reinforced choices (alternative choices or omissions) within the first 5 trials of the extinction phase, the null hypothesis is rejected (p = 0.022, Binomial test), and the behavioral transition is considered abrupt. Otherwise, the behavioral transition is considered smooth.

### Associative network and decision-making model

The model consists of a simple network that associates sensory input with two motor outputs (triangles), one each for the left (L) and right (R) responses (Fig. 5). Binary sensory units (ovals) signal the presence or absence of a specific stimulus (including the context) with a 1 or 0, respectively. These sensory units provide excitatory and inhibitory input to the motor units. Hence, the total synaptic input to the motor units is given by:

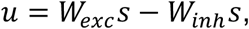

where *u* is a two element vector containing the input to the L and R motor units, *W_exc_* and *W_inh_* are matrices containing the excitatory and inhibitory synaptic weights, respectively, and s is a binary vector specifying the set of stimuli that are present in a given trial. The motor units are rectifying linear units (ReLU), which are driven by the net synaptic input and excitatory noise. Hence, the activation of the motor units is given by:

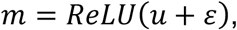

where *m* is a two element vector describing the activation of L and R, and *ε* is a two element vector containing the noise inputs to L and R. These are drawn from two independent uniform distributions on the interval (0, 1). *ReLU*(*x*) = 0 if *x*<0, and *ReLU*(*x*) = *x* if *x*≥0. The behavioral choice corresponds to the motor unit with the highest activation in the presence of a given stimulus and context:

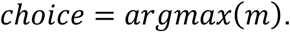

If the total input (synaptic plus noise) to both motor units are equal or lower than 0, no response is selected, resulting in a choice omission.

If a reward is delivered upon responding, excitatory connections between the active sensory units and the responding motor unit are reinforced. Otherwise, excitatory connections remain unchanged. Conversely, when a reward is not delivered, inhibitory connections between the active sensory units and the responding motor unit are reinforced. Otherwise, inhibitory connections remain unchanged. The value of the synapses (i.e. associative strengths) are updated according to:

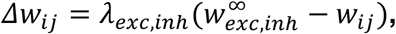

where *Δw_ij_* is the increase of the synapse connecting input *i* with motor unit *j*, *λ_exc,inh_* corresponds to the learning rate of excitation or inhibition, and 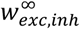 is the maximum possible value that excitatory or inhibitory weights can reach, i.e., their respective saturation values (Bush and Mosteller 1951; Mackintosh 1975; Pearce and Hall 1980; Rescorla and Wagner 1972; Schmajuk and Moore 1985).

For simplicity, all the synapses in the pigeon-models had the same parameters: The learning rates of all the excitatory connections to motor units (λ_e_) were set to 0.02. The learning rate of inhibitory connections (λ_i_) was set either to 0.005, 0.01 or 0.02. This was a way to test the effect of three different ratios of inhibitory/excitatory learning rates (λ_i_/λ_e_ in Fig. 6B and C). All synaptic weights saturated at a value of 20. All synaptic weights were initialized to zero, except for the excitatory weights corresponding to the familiar control stimuli, which were initialized at their saturation value to model the extensive experience that animals had with the familiar stimuli prior to training on the novel stimuli and their near perfect performance. As the familiar stimuli are never subject to extinction, their corresponding inhibitory weights were initialized to zero. The results depicted in Figure S3 correspond to a simplified version of the protocol, where only one extinction context was used, and the familiar stimuli were omitted, to demonstrate the principles of operation of the model more clearly. All other simulations included the two familiar stimuli and two extinction contexts, as described in Figure 1A.

## Results

### Session-to-session variability of choice behavior

In the following, we analyze the responses to the extinction stimulus during the extinction and renewal-test phases, and their dependence on the number of experimental sessions to which an animal was exposed. In particular, we focus on the return of the learned choice during the renewal-test phase, and on the expression of the alternative choice (i.e., pecks on the other available button; a stimulus-response mapping for which the pigeon was not rewarded in the acquisition phase) during the extinction phase (Fig. 2). Figure 2A1 shows the average proportion of learned choices emitted in response to the extinction stimulus as a function of session number. For any given session, the responses emitted during the second half of acquisition (blue), the second half of extinction (red), and renewal-test (black) are shown. During the acquisition phase, animals exhibited a strong preference for the learned choice, which was not significantly correlated with the session number (r = −0.27; p = 0.29). This indicates that animals successfully learned the correct mapping between the novel stimulus and the pecking side repeatedly. During the 2nd half of the extinction phase, on the other hand, the low expression of learned choices show that animals successfully extinguished the learned association in context B. Again, this behavior was also not significantly correlated with session number (r = −0.40; p=0.098). Upon return to context A, the preference for the learned choice reemerged when the extinction stimulus was presented in the absence of reinforcements, which constitutes the ABA renewal effect. Indeed, the proportion of learned choices during the renewal testing phase was significantly higher than during the second half of extinction in 14 out of the 18 session blocks analyzed (p<0.05, z-test; stars in Fig. 2A1). Interestingly, renewal in response to the extinction stimulus changed across repeated sessions of the ABA-renewal paradigm. Across animals, both the average preference for the learned choice (Fig. 2A1, black trace), and the prevalence of these responses (Fig. 2A1, gray bars) were strongly correlated with session number (r = −0.78; p = 0.00024 and r = −0.76; p = 0.00028, respectively). We note that in individual pigeons, the strength of the renewal effect was variable (Fig. 2A1, middle) and intermittent across sessions (Fig. 2A1, bottom). However, the decay of the renewal effect observed across animals was also apparent in individual pigeons. Four animals showed a significant negative correlation between preference and session number (Fig. S1B). The remaining animals also showed negative correlations (range: −0.05 to −0.7) but they were not significant (p > 0.065). Furthermore, the largest response rates for the learned choice during renewal-test occurred within the first four sessions for every individual pigeon (Fig. 2A1, middle).

**Figure 2.**
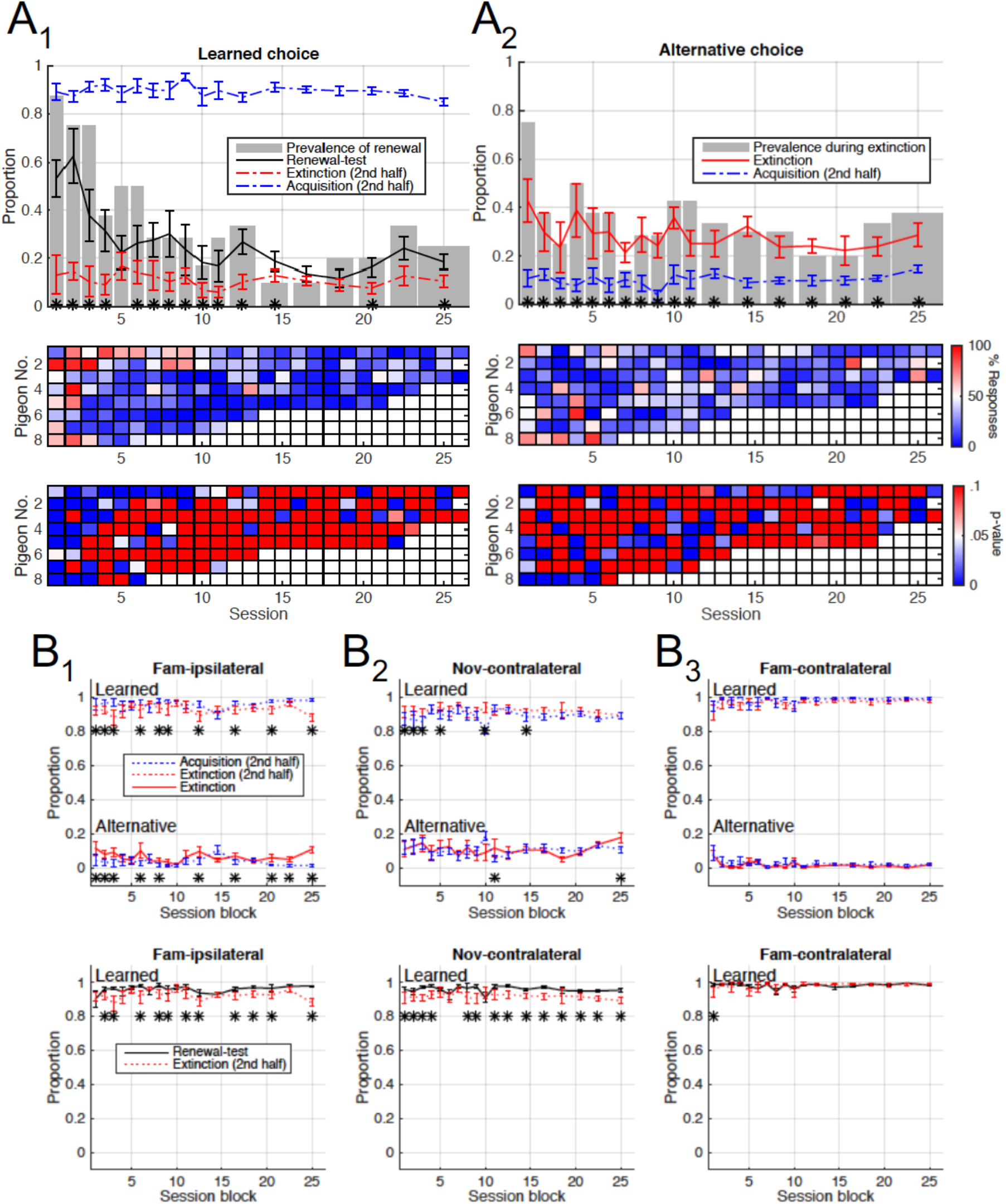
Session-dependent variability of choice behavior. (A) Top: Average response rate across pigeons (lines) and proportion of pigeons expressing a significant number of responses (bars) as a function of session block. Data from sessions >11 were grouped in blocks of 2 to 3 sessions. Bar widths indicate the size of each block. Stars denote a significant difference (p<0.05; z-test) between adjacent traces for each block. Middle: Response rates (color coded) for the learned choice during renewal-test (A1) and for the alternative choice during extinction (A2) for all pigeons and sessions. Bottom: p-values (color coded) of the respective response rates shown in middle for all pigeons and sessions. During renewal-test, both the preference and prevalence of the learned choice (black trace and gray bars in A1, respectively) decay with session number. During extinction (A2), the proportion of pigeons exhibiting a preference for the alternative choice is significantly higher during the first session. (B) Average rate of learned and alternative responses across pigeons for the control stimuli as a function of session block. Responses to the familiar stimulus mapped to the same side as the extinction stimulus (B1), and to the novel and familiar stimuli mapped to the contralateral side (B2 and B3, respectively) are compared across phases (top: Extinction versus Acquisition; bottom: Renewal-test versus extinction).

**Figure 3.**
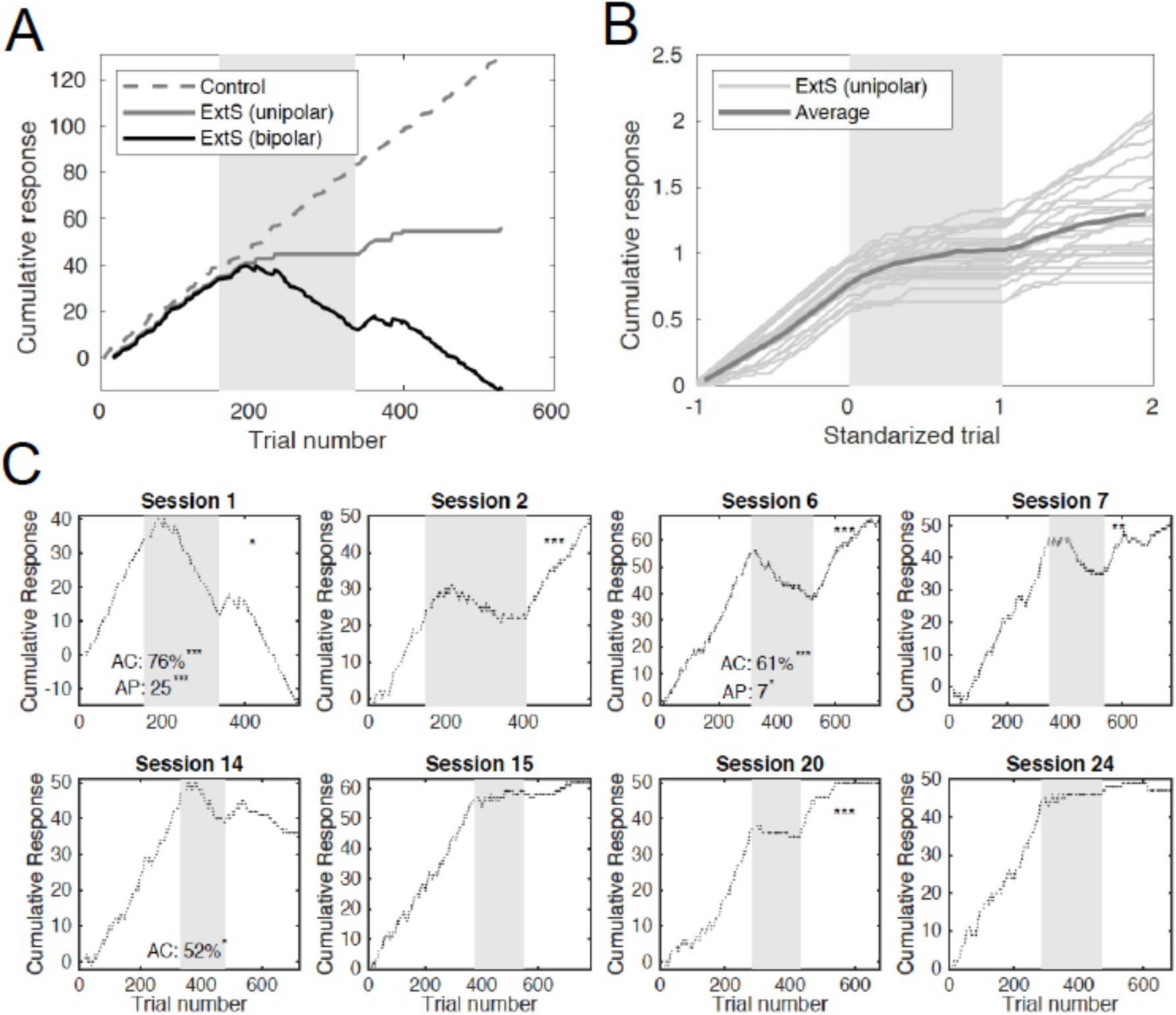
Visualization of choice behavior in single sessions. (A) Single session learning curve obtained using two different choice codings. The trial choices (learned, omission, alternative), can be encoded in a unipolar (1, 0, 0) or bipolar (−1,0,+1) fashion. Unipolar coding (solid gray trace) shows the decay of the learned response in the extinction phase (gray area) and the renewal of the learned response upon return to context A. The bipolar coding (black trace) uncovers the choice behavior during the extinction phase, where a negative slope shows preference for the alternative choice over omissions in the extinction and renewal-test phase. Responses to control stimuli (dashed gray trace) in interleaved trials remained consistent throughout the session. (B) Cumulative record of unipolar-encoded curves for all sessions (thin traces) obtained from one animal along with the grand average (thick trace). Trial numbers are standardized for visualization and averaging (Acquisition: [-1 0); Extinction: [0 1); Renewal: [1 2)). (C) Bipolar-encoded cumulative learning curves for a sample of eight sessions from one animal. Proportion of alternative-choice (AC) responses and alternative persistence (AP; number of consecutive trials yielding an alternative choice) during extinction (gray area) are shown when significant (p < 0.05). Stars on the top-right corner of panels mark the significance of the renewal effect.

**Figure 4.**
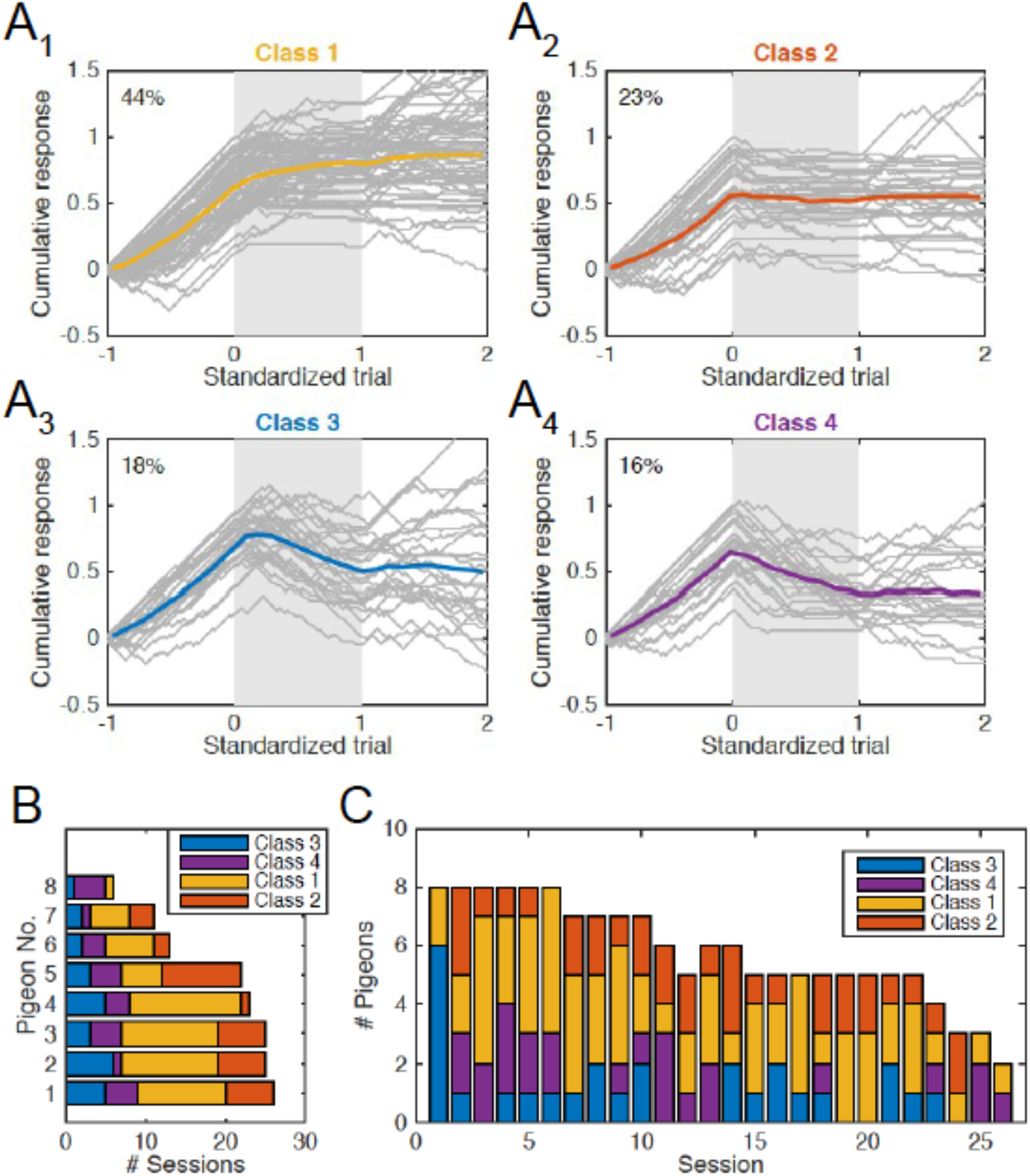
Variability of behavior during extinction. (A) Standardized learning curves (gray traces) corresponding to 151 behavioral sessions obtained from 8 animals. For all sessions, −1 represents the onset of acquisition, 0 the onset of extinction, 1 the onset of the renewal test and 2 the end of the experiment. Curves were classified according to their mode of transition from the acquisition to the extinction phase (smooth vs. abrupt) and their expression of alternative choices during the extinction phase. A1: smooth transition and no alternative choice; A2: abrupt transition and no alternative choice. A3: smooth transition and alternative choice. A4: abrupt transition and alternative choice. Number at the top left corner of each panel indicates the proportion of learning curves that fall into the respective class. (B) Number of learning curves (# sessions) that fall into each of the four classes for each animal. Individual pigeons do not exhibit a clear bias for a particular type of learning curve. (C) Number of animals expressing each class of learning curve as a function of session number. During the first session, all pigeons exhibited smooth transitions at the onset of context B (Classes 1 or 3). Abrupt transitions (Classes 2 or 4) emerged exclusively after the first session.

**Figure 5.**
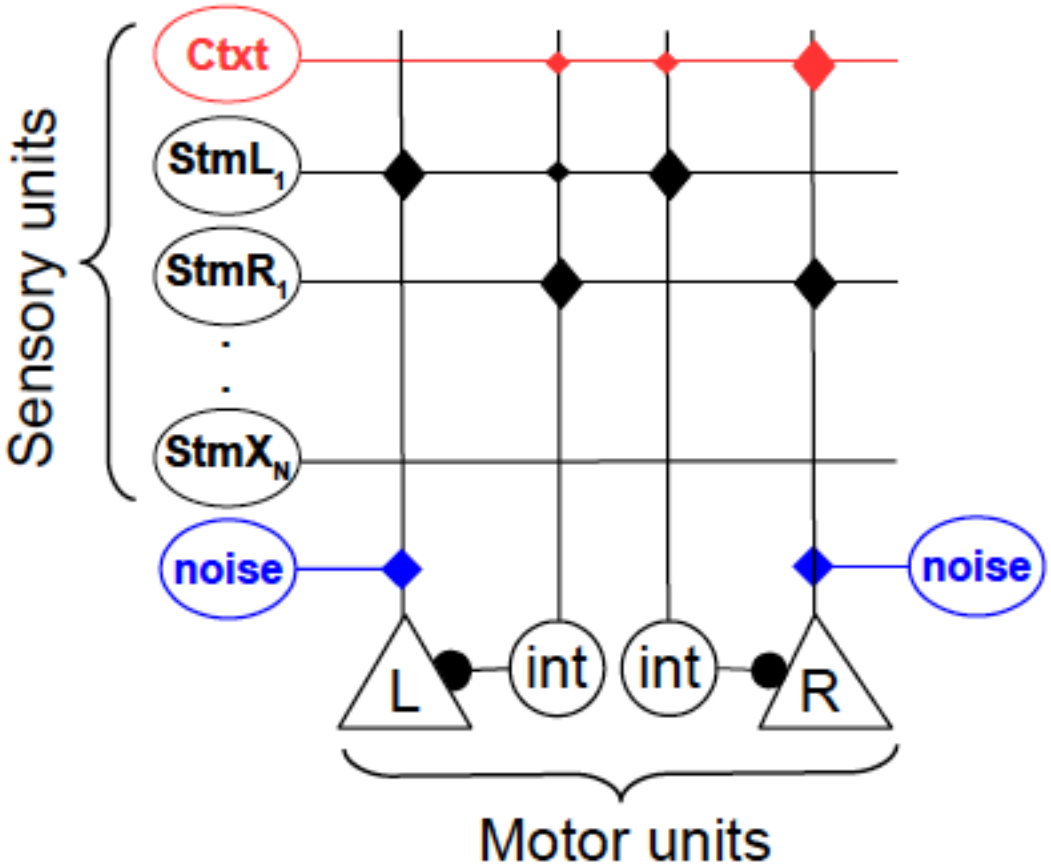
Schematic of associative network model. Sensory units (ovals) can establish excitatory associations directly with motor units (triangles) mediating the left and right responses, or inhibitory associations via interneurons (circles). Context is treated as another simultaneously presented stimulus (red oval; only one context is shown). Motor units also receive excitatory noise (blue ovals). Synapses (diamonds) mediating excitatory and inhibitory associations are reinforced every time a reward is delivered, or not, respectively (diamond size denotes acquired synaptic strengths due to learning). The motor unit receiving the strongest net input generates the respective response. If both units are inhibited below threshold, an omission ensues.

**Figure 6.**
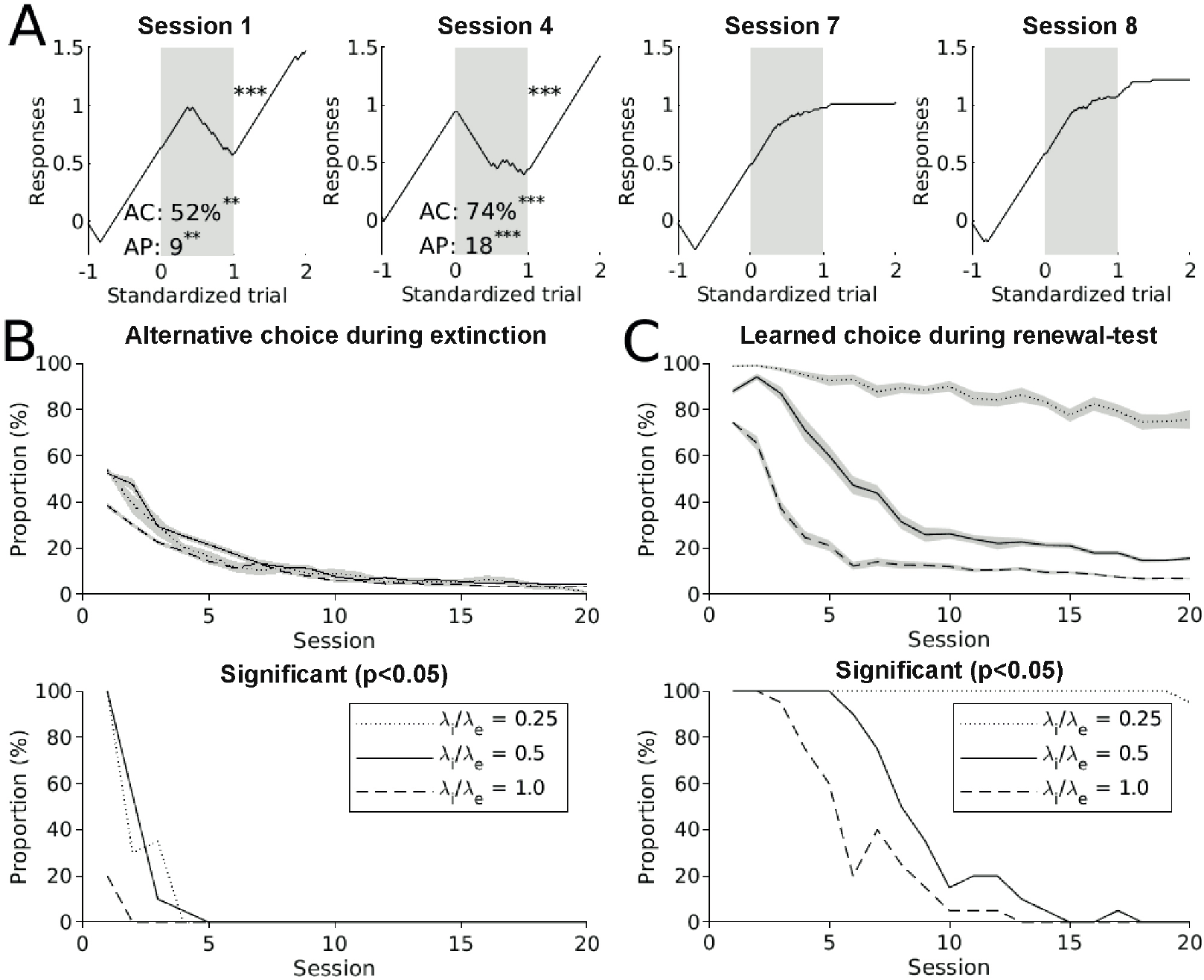
Associative learning accounts for several features observed in the behavior of pigeons in extinction learning and renewal. (A) Sample sessions obtained from one pigeon-model, showing strong preference for the alternative choice (sessions 1 and 4), an abrupt transition at the onset of the extinction phase (session 4) and decay of the renewal effect (sessions 7 and 8). (B) Expression of alternative choice during extinction as proportion of emitted choices (top) and proportion of pigeon-models emitting a significant number of alternative choices (bottom). (C) Prevalence of the renewal effect expressed as proportion of emitted learned choices (top) and proportion of pigeon-models emitting a significant number of learned choices (bottom) during the renewal-test phase. Model results were obtained from a batch of 20 pigeon-models subjected to 20 sessions with randomly selected contexts and extinction stimuli. Simulations were run using three different learning rates for inhibitory connections (0.005, 0.01 and 0.02).

Figure 2A2 shows the proportion of alternative choices emitted in response to the extinction stimulus as a function of session number. During the extinction phase (red trace), the average proportion of alternative choices across animals was significantly higher than during the second half of the acquisition phase (blue trace) for every session block (p<0.05; z-test; stars in Fig. 2A2). In individual pigeons, the expression of alternative choices was also variable (Fig. 2A2, middle) and intermittent (Fig. 2A2, bottom) across sessions. All in all, nearly one-third of the individual sessions exhibited a significant preference for the alternative choice during extinction (Fig. 2A2, bottom). Remarkably, most of these sessions (90%) exhibited chains of 5 to 25 consecutive trials (median: 7 trials) where the animals opted persistently for the alternative choice, further indicating that these responses are not random occurrences. We refer to the length of these chains as ‘alternative persistence’ (AP in Fig. 3C). However, the preference for the alternative choice during extinction was not significantly correlated with session number (r= −0.44; p = 0.070), except for two individual animals (Fig. S1A). In spite of this weak correlation to session number, we nevertheless observed that the prevalence of the alternative choice was not evenly distributed across sessions (Fig. 2A2, gray bars): The proportion of animals expressing this type of responses during the first session (76%) was significantly larger than the proportion found in all the remaining sessions combined (32%; p = 0.0062; z-test). To confirm this asymmetry, we also compared every other session to all the remaining sessions combined, and found no significant difference in the proportions (p > 0.17; z-test).

In our experiment, extinction of one of the novel stimulus-response pairings could have led to a general bias against the response and/or the context that was associated with this extinction learning. Such biases could be driven, for example, by the fact that during the extinction phase, the response and the context associated with the extinction stimulus are rewarded at a ratio of 1:2 relative to the other options. To control for these generalization effects, we also analyzed the responses in the presence of the novel and familiar control stimuli that were presented in interspersed trials (Fig. 2B). In our experimental design, the responses to one familiar control was mapped to the same response (i.e. peck-side) as the extinction stimulus in a different context (Fam-ipsilateral; Fig. 2B1), whereas the other familiar control was mapped to the opposite peck-side in the same context (Fam-contralateral; Fig. 2B3). This design allowed us to study response-generalization and context-generalization separately. During the extinction period, the proportion of learned choices in response to all three control stimuli remained near or above 90% (red dotted line in Fig. 2B). Responses to Fam-ipsilateral were slightly (but significantly) reduced during extinction when compared to the responses during the acquisition phase in 10 out of the 18 session blocks (p<0.05; z-test; stars in Fig. 2B1, top), but the responses to Fam-contralateral remained unchanged (p>0.05; z-test; Fig. 2B3). So, there was evidence for response-generalization of extinction learning, but not for context-generalization.

A general response bias against the response associated with extinction learning would lead to increased pecking on the other side, which we classified as alternative choice. There indeed was a slight significant increase in the alternative choices in response to the Fam-ipsilateral control in 10 of the 18 blocks (Fig. 2B1, top panel). Furthermore, the response bias should affect the contralateral controls (Fam-contralateral and Nov-contralateral) the opposite way, i.e., increasing learned choice and decreasing alternative choices. Indeed, for the Nov-contralateral there was a slight significant increase in learned choices during the extinction phase (Figs. 2B2, top panel), which might however also be driven by continued reinforcement. No effects were observed for the learned choice for the Fam-contralateral control, probably due to ceiling effect, nor for the alternative choices (Figs. 2B2, and 2B3, top panel), probably due to the floor effect. In summary, while there was evidence for a general response-bias, the magnitude of this bias evident in the control stimuli was smaller than the increase in alternative choices for the extinction stimulus. Therefore, a general response-bias does not account for the emergence of alternative choices during extinction learning.

Response changes induced through generalization appear to be sensitive to context change from the extinction contexts, B1 or B2, back to the original acquisition context A during the renewal-test phase. While the proportion of learned choices in response to the three control stimuli remained stable near or above 90% in the renewal-test period (black lines in Fig. 2B, bottom panels), the preference for the learned choice in response to Nov-contralateral (black trace in Fig. 2B2) was slightly higher than during the second half of extinction in 14 out of the 18 session blocks (p<0.05; z-test; stars in Fig. 2B2, bottom panel). Importantly, none of the responses to the control stimuli were significantly correlated with session number (p>0.05, Fig. 2B), confirming that the decay of the renewal effect for the extinction stimulus across sessions was not due to a general decay in the engagement of the animals in the task (e.g., due to satiation).

### Visualizing choice-behavior in single sessions

To study the session-to-session changes in choice behavior in more detail, we visualized the entire time-course of learning within individual sessions. We did this by focusing on the cumulative record of successive behavioral responses as a function of trial number (Fig. 3; Gallistel et al. 2004; Leslie et al. 2005). However, we departed from the traditional ‘unipolar’ encoding of the responses, which focuses only on the learned choice by signaling its presence or absence with 1 or 0, respectively. Instead, we encoded each learned choice, alternative choice, or omission as +1, −1 and 0, respectively. Under this bipolar encoding, the slope in the cumulative curve reveals biases towards specific responses (Fig. 3A, black trace): A positive or negative slope indicates a tendency to prefer the learned choice or the alternative choice, respectively. A slope of 0 indicates either a continuous chain of omissions or random mixtures of different responses. In Figure 3A, for example, the bipolar encoding (black trace) reveals that the extinction phase (gray area) is characterized by a persistent selection of the alternative choice. For comparison, the gray traces in Figure 3A and B illustrate how the unipolar encoding conceals the choice behavior. By focusing on a target response, the unipolar encoding can create the impression that the behavior observed in a grand average across sessions is representative of single sessions (Fig. 3B). Furthermore, the average unipolar learning curve looks similar to what one expects to find in ABA renewal experiments that use very different experimental designs from ours (e.g. Bouton et al. 2011; Gottfried and Dolan 2004; Lissek et al. 2015). Since our focus is on choice behavior, in the following, we use bipolar learning curves to study the variability of behavior across multiple sessions (Fig. 3).

### Diversity of learning curves across sessions

From session to session, individual animals exhibited a rich repertoire of learning curves. As an example, Figure 3C shows eight sessions obtained from the same animal. In these examples, the behavior during the extinction phase (gray area) is characterized by random responses (e.g., sessions 2, 7 and 15) or omissions (e.g., sessions 20 and 24). In some cases, however, the learning curves developed negative slopes (e.g., in sessions 1, 6 and 14), revealing the preference for the alternative choice (AC in Fig. 3C) described in the previous analysis (Fig. 2A2). In the same way, the renewal-test phase was characterized by a variable re-emergence of the learned response (stars on top right of panels show significance of the effect; see also Fig. 2A1, bottom). Upon return to context A, some sessions were characterized by a few learned responses before re-extinction took place (e.g., sessions 1, 7 and 20), or exhibited no significant renewal (e.g., sessions 15 and 24; see also Fig. 2A1, bottom). In some cases, animals persisted on the learned choice for most of the renewal-test phase (e.g., sessions 2 and 6; see also Fig. 2A1, middle).

Another puzzling aspect uncovered by the bipolar encoding were the abrupt transitions upon the onset of the extinction phase. According to a widespread view of extinction learning, as animals experience a withdrawal of reinforcers upon the onset of context B, they initially persist on the previously reinforced choice for several trials before gradually changing their behavior towards omissions (Nevin 2012; Podlesnik and Shahan 2009, 2010). Indeed, several learning curves in our data were characterized by these smooth transitions (e.g., sessions 1 and 2 in Fig. 3C). In some sessions, however, the change in behavior seemed to occur abruptly upon the onset of context B (e.g., sessions 6, 7, 20 and 24 in Fig. 3C). These changes were apparently driven by the switch from the acquisition to the extinction context. To assess the presence of this peculiar behavior, we performed a categorical distinction based on the behavior within the first 5 trials of the extinction phase. Namely, we considered a transition abrupt if a pigeon emitted at least 3 non-reinforced choices (alternative choices or omissions) within the first 5 extinction-trials, which rejects the hypothesis that animals continued to perform as required to end the acquisition phase (p = 0.022, Binomial test; see Materials and Methods). According to this criterion, 39% of the sessions exhibited such abrupt transitions.

### Four extinction types and their session to session variability

So far, we have identified two remarkable features of extinction learning, namely, the preference for the alternative choice, and abrupt transitions of behavior upon the onset of the extinction phase. Based on the presence or absence of these features, we arranged the learning curves in four classes, and then assessed the prevalence of these classes across pigeons and sessions (Fig. 4) to see whether there are systematic preferences for a certain type of learning curve. Figure 4A shows a two-by-two matrix of panels where the left and right columns correspond to smooth and abrupt transitions, respectively. And the top and bottom rows to the absence and presence of alternative choices, respectively. Each panel contains an overlay of all the learning curves (gray traces) belonging to the respective class, and their average trace (colored line). Thus, class 1 corresponds to the canonical learning curves, which accounted for 44% of the total sessions (Fig. 4A1). The majority of the learning curves (56%) deviated from the canonical one in at least one major aspect: either because animals exhibited abrupt transitions at the onset of context B (class 2), favored the alternative choice over omissions during extinction (class 3), or both (class 4, Fig. 4A4). Next, we analyzed the way in which the learning curve types described in Figure 4A were distributed across pigeons and sessions (Fig. 4B and C, respectively). Individually, pigeons did not consistently express a particular class of learning curve. In fact, all pigeons, except for one (for which only 6 sessions were available), exhibited all four learning curves classes (Fig. 4B). To get insights on the effects of re-testing the animals repeatedly, we also analyzed the session-to-session changes in the distribution of learning curve classes (Fig. 4C). Curves with smooth transitions (classes 1 and 3) were more prevalent during the first session, and occurred less frequently in later sessions. This last observation, however, was based on a categorical variable. To provide further quantitative support for this finding, we used the method proposed by Gallistel et al. (2004) to determine the first change point of the learning curves during extinction. We found that the change of behavior occurred significantly later in the first session as compared to the subsequent sessions (p = 0.023; KS-Test, Fig. S2).

### Can associative learning account for the observed behavioral complexity?

So far, we have reported three key findings in our behavioral data: (1) Smooth transitions of behavior at the onset of context B are more prevalent during the first exposure to the extinction task than in later sessions. (2) During extinction, pigeons express a preference for the alternative choice intermittently across sessions, most prominently in the first session. And (3), the renewal effect appears intermittently in individual pigeons, but shows an overall decay as sessions progress. To study the mechanisms underlying these phenomena, we implemented a parsimonious model aimed at capturing the associative aspect of the task (Fig. 5). This model embodies two fundamental principles of associative learning. First, representations of the present stimuli can freely establish both excitatory and inhibitory associations with motor actions. Second, these associations are modulated by reinforcement contingencies. Additionally, we treat the context as just another stimulus that can establish direct inhibitory and excitatory associations with specific motor actions, in accordance with recent studies (Bernal-Gamboa et al. 2018; Bouton 2019; Nieto et al. 2017; Todd 2013; Todd et al. 2014).

Figure 5 summarizes the components of the model, which operate as follows: Sensory units (ovals) signal the representation of the context and specific stimuli in working memory. These units can establish direct excitatory connections with the motor units (triangles) mediating the left (L) and right (R) responses. They can also inhibit the motor units via interneurons (circles). Thus, excitatory and inhibitory associations between stimuli and actions are mediated by two independent pathways, as previously suggested (Felsenberg et al. 2018; Ghazizadeh et al. 2012). The decision making is performed by a simple, instantaneous winner-takes-all mechanism, where the motor unit with the highest activation drives the corresponding behavioral response. If both motor units are inhibited below their threshold of activation, no response is selected, resulting in a choice omission. Excitatory connections to motor units are reinforced if a reward is delivered, and remain unchanged otherwise. Conversely, connections onto interneurons are reinforced only if a reward is not delivered, and remain unchanged otherwise. The synaptic weights (i.e., associative strengths) mediating these connections grow asymptotically towards a saturation value, in accordance with standard models of associative learning (Bush and Mosteller 1951; Mackintosh 1975; Rescorla and Wagner 1972).

### Associative learning accounts for the features observed in the behavioral data

We subjected a population of 20 pigeon-models to a sequence of 20 training sessions following the same experimental protocol depicted in Figure 1A, where the extinction context and extinction stimulus were selected randomly. For simplicity, all the synapses in the pigeon-models had the same parameters (see Methods for details). We run this simulated experiment using three different values for the learning rate of inhibition (λ_i_), which allowed us to test the effect of three different ratios of inhibitory/excitatory learning rates (λ_i_/λ_e_ in Fig. 6B and C). Figure 6A shows a sample of four learning curves obtained from one of the pigeon-models. These curves exhibit three features we uncovered in the pigeon behavior: preference for alternative choices during extinction (sessions 1 and 4), abrupt transitions of behavior upon onset of context B (session 4), and absence of the renewal effect (sessions 7 and 8). Like the pigeons in our study, the model shows a preference for the alternative choice during extinction, which rapidly declines as a function of session number (Fig. 6B, top). The proportion of pigeon-models emitting a significant number of alternative responses (Fig. 6B, bottom) resembles the findings in our experimental data (see Fig. 2A). Across the set of parameters tested, the significant expression of alternative choices was limited to the first few sessions. Also qualitatively reproducing our observations in pigeons, the expression of the renewal effect declined as a function of session number, as evidenced by both the average preference for the learned choices during the renewal-test phase (Fig. 6C, top), and in the proportion of pigeon-models emitting a significant number of learned choices (Fig. 6C, bottom). Since no parameter was adjusted between sessions, the variability in behavior stemmed solely from the history of learning. In particular, due to the associations between the context and specific responses (Bouton 2019; Leitenberg et al. 1970; Todd 2013; Todd et al. 2014; Winterbauer and Bouton 2010), which are carried over from session to session.

The precise mechanisms underlying this variability are explained in detail in the supplementary information (see Fig. S3 and accompanying text), where we used a simplified version of the protocol consisting of only two stimuli and one context. In a nutshell, the model embodies three core assumptions: (1) that context and stimuli can both drive and inhibit a given response in parallel, (2) that drives are upmodulated by the presence of reinforcements, whereas inhibitions are increased when reinforcements are absent, and (3) the response with the strongest net drive is expressed. As this model is subject to repeated sessions of learning and extinction, a gradual build-up of drivers and inhibitors within and throughout sessions ensues. Here, we aim at describing an extinction scenario where, under the same context, some previously learned stimulus-response mappings continue to be rewarded (our control stimuli) but one is not (our extinction stimulus). Since stimuli are session-unique, but context and responses recur in multiple sessions, previously established context-response drives and inhibitions can be carried over subsequent sessions, and thereby bias the responses to new stimuli presented in the same context. Thus, once the first session is completed, two processes with different time-scales will compete in the upcoming sessions: A fast process triggered by the previously established context-response mappings, and a slow process of learning the new stimulus-response mapping. The fast-process can drive an abrupt change of behavior upon a context switch, but the larger the number of trials, the stronger the effect of the slow cumulative process becomes. More specifically, within extinction sessions, negative associations between context B and the learned choice are established due to the absence of reinforcers. Conversely, positive associations between context B and the contralateral response are also established during the interspersed trials where the control stimuli are presented, and their learned responses still rewarded. Together, these associations act synergistically to tilt the balance in favor of the alternative choice. As sessions progress, the associative strengths maintaining this imbalance saturate quickly, favoring the expression of alternative choices during the first session. The prevalence of smooth transitions during the first session can be explained as follows: Upon the first exposure to the extinction phase, negative associations between the context and specific motor responses require several trials to build up (e.g., Fig 3SC1), but once established, they can exert an effect on the behavior in later sessions (e.g., Fig. 3SC2–4). This inheritance of context-response associations can lead to abrupt transitions of behavior upon switching to the extinction context, which can only occur in sessions after the first exposure to a context-dependent extinction phase. The renewal effect, on the other hand, occurs due to the release of the inhibition exerted by context B on a specific response (Bouton 2019; Todd 2013; Todd et al. 2014; Trask et al. 2017). After many sessions of training, these negative associations between context and responses reach their saturation value. As a consequence, the learned response can no longer be rescued by the release of context-inhibition. Thus, the decay of the renewal effect with sessions is a natural consequence of the existence of an asymptote of conditioning; a property that is ubiquitous across models of associative learning (Bush and Mosteller 1951; Mackintosh 1975; Pearce and Hall 1980; Rescorla and Wagner 1972; Schmajuk and Moore 1985).

Although our model was able to capture the main features observed in the responses to the extinction stimulus, it did not reproduce the slight but significant biases observed in the control stimuli (see Fig. 2B). In the model, all controls reached perfect performance during acquisition (i.e. 100% conditioned choice), and remained at the same level during extinction and renewal across sessions (not shown). Consistently, alternative choices were exclusively expressed in response to the extinction stimulus. This is presumably due to the fact that the excitatory connections between the familiar stimuli and their respective responses were initialized at their saturation value (see Materials and Methods for justification). Since in our simulations, inhibitory connections saturate at the same level as excitatory ones, it is not possible to tilt de balance in favour of an alternative choice.

## Discussion

We have analyzed the choice behavior of pigeons subject to multiple sessions of a discrimination learning task in context A, extinction in context B, and a return to context A to test for the return of the learned response. By focusing on the expression of the learned and alternative choices from individual animals in single sessions, as well as on the corresponding learning curves, we uncovered a rich choice variability within and across sessions during the extinction and renewal-test phases: (1) Upon the onset of the extinction phase, pigeons tended to persist on the learned choice mostly during the first session, whereas abrupt transitions of behavior emerged exclusively in later sessions. (2) During extinction, the expression of alternative choices in individual pigeons appeared intermittently from session to session, most prevalently in the first session. And (3), the renewal effect in individual pigeons was also intermittent and variable, but its strength and prevalence decayed as sessions progressed. To reveal potential mechanisms of this behavioral variability, we used a computational model to show that associative learning, in combination with an instantaneous winner-takes-all decision process, can reproduce the aforementioned features observed in the data.

### Experimental paradigm and methodology

There are several aspects in our paradigm that differ from previous studies of repeated extinction. These differences allowed us to focus both on the expression of alternative behaviors during extinction and the dynamics of the renewal-effect. First, our discrimination-learning task comprises more than one operant response (learned and alternative choices as well as choice omissions), in contrast to previous studies that used a single response key (e.g. Anger & Anger, 1976; Bai and Podlesnik, 2017), lever-press responses (Samson et al., 2001) or head entry responses (Guilhardi et al., 2006). This methodological difference granted the expression of variable behaviors during repeated extinction and renewal. Furthermore, previous studies in which alternative operant responses are available, often do not analyze their expression (e.g., André et. al., 2015). Another methodological aspect that differs from previous studies is our highly standardized procedure of within-session acquisition, extinction and renewal. Here, imposed fixed behavioral criteria that have to be met in order to transition from phase to phase. In this way, we minimized variations in the acquisition procedure (e.g., in contrast to the repeated re-acquisitions in the case of Anger & Anger, 1976), changes in reinforcement schedules within the experimental phases (e.g., Gershman et al. 2010) or differences in the acquisition/extinction intervals. Thus, our method allows a direct comparison of subsequent sessions, including the renewal-effect, which goes beyond the previous reports on repeated extinction. Yet another aspect that distinguishes our paradigm from previous ones is the way we switch contexts by changing the house-lights color and the auditory feedback in response to the key pressings. Although context often refers to the enclosure where the experiment takes place, there is considerable experimental evidence that, to observe the effects of contextual changes, it is not required that a complex environment is present for a long period of time and then changed radically. Instead, it is sufficient that a salient fraction of contextual stimuli is present during the presentation of the nominal stimulus (for an explicit proof, see Lachnit 1986; Üngör and Lachnit 2006; for successful application, see Uengoer et al. 2013).

Finally, it is possible that some aspects of the behavioural variability reported here (e.g., the intermittence of the renewal effect) might have been concealed by averaging in previous studies (André et al. 2015; Cheng and Sabes, 2006; Gershman et al. 2010; Larrauri and Schmajuk 2008; Lissek et al. 2015; Todd 2013; Trask et al. 2017; Wirth et al. 2003). The issue of averaging is also relevant for computational models, as they usually assume that the properties of the average curve are representative of those of individual curves. This assumption prevents models from giving a comprehensive account of the phenomenon of extinction learning, in particular with regard to the expression of choice behavior and its variability across subjects and sessions.

### Function and mechanisms of alternative responses

The emergence of apparently purposeless behavior is puzzling. If one particular response is abandoned due to the lack of reinforcers, and an alternative is picked as a result, why would an animal persist on an unrewarded alternative over omissions? Evolutionary theories of foraging have proposed reasons why probing alternative behaviors might pay under reduced reward conditions (Anselme and Güntürkün 2019; Gharib et al. 2004), but these arguments explain the ultimate level, thereby lacking proximate, mechanistic explanations. From the perspective of reinforcement learning, explaining unrewarded alternative choices is difficult, as these responses do not provide any added value relative to an omission, and incurs the cost of the extra energy spent. For instance, the model by Redish et. al, (2007) predicts that extinguishing one action in response to a stimulus results in the suppression of all available actions in response to the same stimulus (Redish et al. 2007). Our computational model, on the other hand, provides a proof-of-principle that associative learning, combined with a winner-takes-all decision-making process, can give rise to exploratory behaviors and persistence on previously unrewarded responses. Thus, we offer a parsimonious, putative mechanism to account for the emergence of exploratory behaviors under altered reinforcement contingencies.

### Associative learning: simple explanations for seemingly complex cognitive processes

Although our model provides a parsimonious explanation of the observed behavior in terms of associative learning, it cannot rule out the involvement of higher-order cognitive functions. Indeed, several of the features observed in the behavior can be interpreted as an expression of animals learning about the structure of the task (Livesey et al. 2019; Zentall et al. 2014), and exploiting that knowledge to maximize reward. For example, it has been suggested that when animals are exposed to several sessions of extinction, they might learn to discriminate the transition to extinction (e.g., Baum 2012; Bullock and Smith, 1953). In our case, animals might have learned to discriminate the transitions between experimental phases signalled by context changes. This could explain, for example, the decay of the renewal effect with session number (Fig. 2A1): As sessions progress, animals might learn that a switch from context B back to context A does not predict a return of the reinforcement contingency of the acquisition phase. Hence, the renewal effect would decay gradually as animals experience more and more sessions with the same ABA structure. Since such putative abstract-rule learning is not perfect, some forgetting or attentional fluctuations might lead to the intermittent reappearance of the renewal effect (Fig. 2A1, bottom), but overall, renewal decays with experience. Along the same lines, the predominance of smooth transitions in the first session, and the appearance of abrupt transitions at later sessions (Fig. 4C) might also reflect some form of abstract-rule learning: In the first session, animals experience a withdrawal of reinforcers when the context is switched to B for the very first time. Not knowing that reward contingency is linked to context, the animals will initially persist on the previously reinforced response for several trials before gradually changing their behavior as a result of the absence of the expected reward. Such behavioral momentum might be reduced in later sessions as animals learn that a change from the acquisition context A to the extinction context B signals a change in the reward contingency. The application of this hypothetical rule might also be subject to attentional fluctuations and other sources of noise, resulting in the observed intermittent pattern of abrupt transitions.

With respect to the expression of alternative choices during extinction, on the other hand, it could be argued that they emerge due to the asymmetry in global rates of reinforcement during our extinction paradigm (one pecking side is reinforced at twice the rate of the other). Under uncertainty (e.g. during extinction trials) animals might simply opt for the response that is most often rewarded, which is the alternative choice for the extinction stimulus. This is particularly the case during the initial phase of the first extinction session, where animals experience for the first time that the previously learned responses to the extinction stimulus are no longer rewarded. At some point during the extinction phase, the animal has collected enough evidence that pecks to the side contralateral to the learned response for the extinction stimulus are more often rewarded than the ipsilateral responses. Therefore, the contralateral response would be the most “reasonable” to go for under uncertainty (the animal is not uncertain in the presence of the other control stimuli, as their response mappings are still reinforced as usual). The fact that alternative choices occur more often during the first sessions than in later ones is consistent with this view. As animals need to be exposed to at least one session in order to “understand” the structure of the task, repeated extinction reduces the uncertainty and, hence, the proportion of alternative choices. While this apparently “rational behavior” is consistent with our data, this interpretation is not necessary, since this pattern of behavioral changes emerges in a simple associative learning model.

Yet another possibility to explain some of our results would be in terms of learned equivalence (Honey and Hall 1989; Zentall et al. 2014). In this framework, pigeons exposed to repeated sessions of our experimental paradigm could learn to group the stimuli under two context-sensitive categories, namely ‘Peck-Left’ and ‘Peck-Right’. During extinction, alternative responses would emerge from a re-categorization of the extinction stimulus. After a few sessions, one could hypothesize that animals learn that the extinction context signals this re-categorization, giving rise to abrupt transitions of behavior upon the onset of context B. Based on our data, we cannot rule out that these hypothetical behavioral strategies drive the observed complex behaviors during extinction and renewal-test. However, in accordance with Morgan’s Canon (Morgan 1903), we favor the parsimonious explanation of our results provided by our simple associative learning model over an interpretation in terms of higher-order cognition.

### Preference for alternative choices and resurgence

The expression of alternative choices in our experiment could be related to the phenomenon of resurgence, wherein a previously reinforced and then extinguished response reappears during a period of extinction for a subsequently learned response (Leitenberg et al. 1970; Winterbauer and Bouton 2010). Thus, if we consider that an alternative choice in a given session might correspond to a response that has been reinforced and extinguished in a previous session, this previously extinguished response might “resurge” in the current session as an alternative choice once the reinforced choice is extinguished. However, in our results, alternative choices occur prominently already in the first extinction session, before any responses had been extinguished. In addition, resurgence cannot account for any of our other findings, namely, abrupt transitions, lack of renewal, and the dynamics of these behaviors.

### The role of context in extinction and renewal

A key aspect of our model is that the context can directly establish excitatory and inhibitory associations with specific responses (Bouton 2019; Rescorla 1993, 1997; Todd 2013; Todd et al. 2014). This feature was critical for our model’s ability to account for the data. In particular, alternative responses emerged in the model, because it learned an excitatory association between the extinction context and the alternative choice either during an earlier phase or during the extinction phase when the response to the control stimuli are rewarded. However, it has been suggested that context can also establish associations with the representations of the outcome (Baker et al. 1991; Pearce and Hall 1979), or modulate the association between response and outcome (occasion setting hypothesis) (Trask and Bouton 2014). Since these hypotheses are not mutually exclusive, our model alone cannot rule out a scenario where context exerts a direct or modulatory influence over the representation of the outcome. We also did not consider associations between context and stimuli. However, it is hard to see how a context-cue association could drive any variability across sessions in our behavioral paradigm, because the novel stimuli are session-unique. Therefore, previously established associations between the context and a novel stimulus presented in a previous session plays no role in subsequent sessions because that stimulus does not repeat.

### Scope of the model

The purpose of our model was to test whether highly variable individual behaviors can emerge from simple associative learning. Therefore, we included several elements from existing associative learning models, but also excluded components that could be attributed to higher cognitive functions in those models. Specifically, the law governing the update of associative strengths in each unit of our model is the same as the one used by Bush and Mosteller (1951), but with the associability term (or salience) set to 1. Other models have used this term to describe the selective attention certain stimuli become due to their relative predictive power of desired/undesired outcomes (Bush and Mosteller 1951; Mackintosh 1975; Pearce and Hall 1980; Rescorla and Wagner 1972; Schmajuk and Moore 1985). In some of these models, this term is updated from trial to trial, thereby modulating the learning rates dynamically within single sessions (Mackintosh 1975; Pearce and Hall 1980; Schmajuk and Moore 1985) (for empirical evidence for learning-induced changes in selective attention, see Feldmann-Wüstefeld et al. 2015; Lucke et al. 2013; Koenig et al. 2017a, 2017b). Curiously, our model exhibits extinction learning at varying speeds (see Figs. 6A and S3C), even though it lacks a term modulating the learning rates; a phenomenon that could be otherwise attributed, for example, to attentional variations due to reward expectancy.

### Possible clinical implications

The present results have specific implications for the treatment of pathological behaviors (Podlesnik et al. 2017). The higher prevalence of alternative choices during the first session in the present experiment indicates that treatments aimed at extinguishing problem behavior and promoting alternative, appropriate behavior should make particular use of the early stage of extinction treatment to reinforce the more desirable alternative responses that might emerge (Antonitis 1951; Fuller 1949; Grow et al. 2008; Lattal and Lattal 2012; Tinsley et al. 2002). Moreover, the decay of the renewal effect with session number suggests that repetitions of a treatment involving extinction can be helpful in order to prevent the context-dependent reemergence of undesired behaviors.

In conclusion, we have uncovered a rich variability of behavior in extinction learning and renewal that so far has remained concealed in population averages. Even though these complex behaviors appear to reflect abstract rule learning, we have demonstrated that associative learning can generate similarly complex behavior without resorting to higher-order cognitive processes.

## Author Contributions

Designed research: JRD, JP, RP, MU, HL, OG, SC; Performed research: JRD, JP, RP; Contributed new reagents or analytic tools: ZL, TW; Analyzed data: JRD, ZL, SC; Wrote the paper: JRD, JP, RP, MU, HL, OG, SC.

## Acknowledgments

Funded by the Deutsche Forschungsgemeinschaft (DFG, German Research Foundation) – project number 316803389 – SFB 1280, projects F01, A01, A14 and A15.

## Compliance with Ethical Standards

### Funding

This study was funded by the Deutsche Forschungsgemeinschaft (DFG, German Research Foundation) – project number 316803389 – SFB 1280, projects F01, A01, A14 and A15.

### Ethical approval

All animals were treated in accordance with the German guidelines for the care and use of animals in science. The experimental procedures were approved by a national ethics committee of the State of North Rhine-Westphalia, Germany and were in agreement with the European Communities Council Directive 86/609/EEC concerning the care and use of animals for experimental purposes.

