## Supplementary Information for "Emergence of complex dynamics of choice due to repeated exposures to extinction learning"

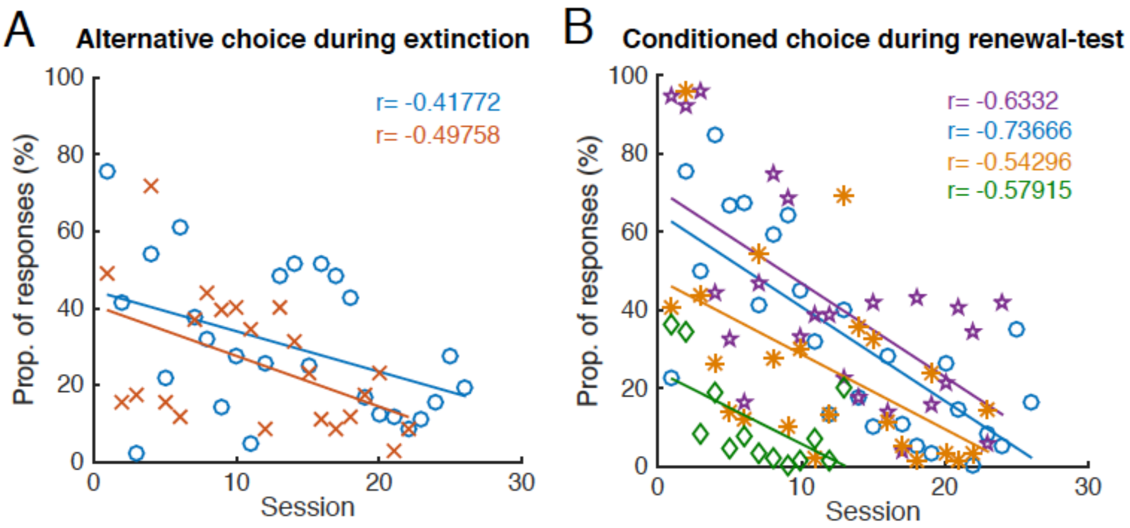


**Figure S1.** *Preference for target response as a function of session number.* (A) Preference for alternative choice during extinction. (B) Preference for conditioned choice during renewal. Only pigeons showing significant correlations are shown (symbols). Correlations for the corresponding linear fits (colored lines) are indicated in the top corner of the panel (colored text).

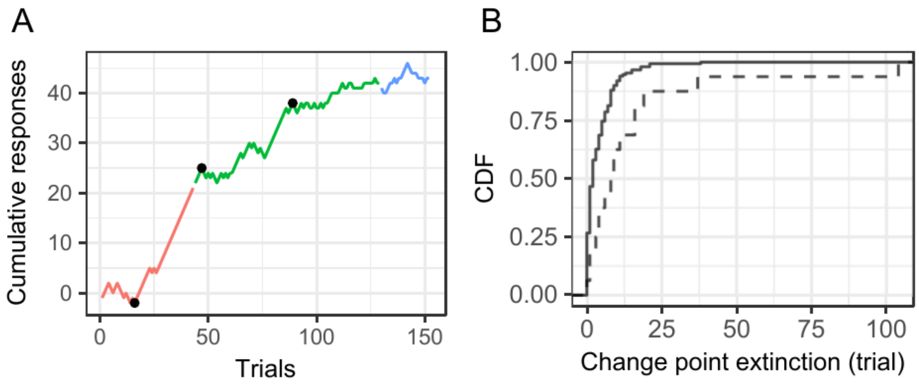


**Figure S2.** *Change point analysis reveals slower extinction during the first session*. (A) Cumulative responses to the extinction stimulus as a function of extinction trial number (only extinction trials are enumerated). Phase is color-coded (red: acquisition; green: extinction; blue: renewal-test). Black dots show the change-points as identified by the change point analysis. (B) Cumulative distribution of the extinction trial number, at which the first change point occurs during the extinction phase (dashed: first session; solid: remaining sessions). The change point occurs earlier in later sessions.

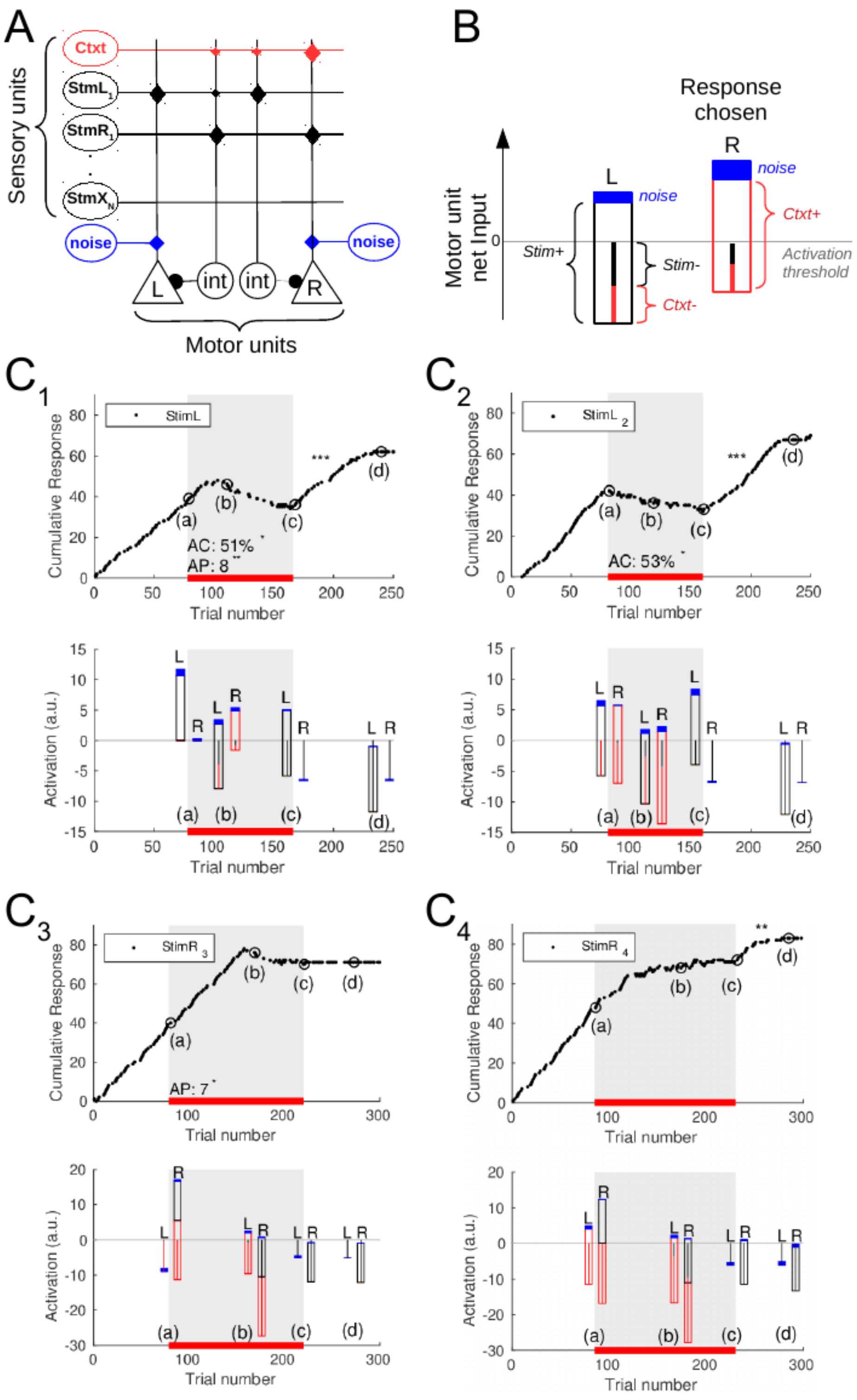


**Figure S3.** *Associative learning can drive complex choice-behavior during extinction and renewal.* (A) Schematic of associative network model. Sensory units (ovals) can establish excitatory associations directly with motor units (triangles) mediating the left and right responses, or inhibitory associations via interneurons (circles). Synapses (diamonds) mediating excitatory and inhibitory associations are reinforced every time a reward is delivered, or not, respectively (diamond size denotes acquired synaptic strengths due to learning). Motor units also receive excitatory noise. (B) Schematic of how the composition of the motor unit input activity is depicted: Inhibition (indicated by vertical lines), excitation (indicated by open bars), and excitatory noise (indicated by solid blue bars). Hence, the net activation is indicated by the top of the bar. The unit with the highest net activation triggers the corresponding behavioral response. If the net activation of both motor units remains below the threshold, a choice omission ensues. (C) Model responses to four consecutive sessions of a simplified version of the task performed by the pigeons. Only one extinction context was used, and control familiar stimuli were excluded (top). Cumulative responses for the extinction stimulus. The proportion of alternative choices (AC) and longest chain of successive alternative choices (AP: Alternative persistence) during extinction are shown. (C1) Preference for the alternative choice during extinction. (C2) Abrupt transition upon onset of extinction context. (C3) Absence of renewal. (C4) Reappearance of renewal. Note that the variable dynamics of extinction emerged due to remnant context-response associations from previous sessions. (bottom) Input composition of the left and right motor-output units (L and R, respectively) in response to the extinction stimulus (black) and context (red). The contribution of each component to the activation is coded as shown in B. Activity is sampled at the onset of the extinction phase (a), during extinction (b), at the onset of the renewal-test phase (c), and at the end of the renewal-test phase (d); see corresponding circles on top. The learning rates of all connections were set to 0.02, and all synaptic weights saturated at a value of 20 (see Materials and Methods). The number of trials in these model sessions are smaller than those in the experimental sessions due to the simplified version of the task.

### Supplementary Text

**How associative networks can drive complex behavior.** To illustrate the basic interactions underlying the behavior of the model, we put the model through a simplified version of the protocol used in the behavioral experiments (Fig. S3). Here, only two stimuli were presented instead of four, and only one extinction context was used instead of two. Briefly, during the acquisition phase, responses to the left or right were reinforced when the corresponding stimuli, StimL or StimR, were presented, respectively. At the onset of the extinction phase, the context unit was activated and no rewards were given in the presence of the extinction stimulus (StimL in Fig. S3C1), regardless of the response given. Figure S3C shows the behavior of the model during the first and subsequent three sessions of the task. These four sessions provide an example of how associative learning can generate the complex choice behavior observed in the experimental data. To illustrate the evolution of the associations giving rise to the behavior of the model, we display the excitatory and inhibitory contributions of both context and extinction stimulus to the activity of the left and right motor units (Fig. S3B and Fig. S3C, bottom panels) at specific points of the learning curve (indicated by red circles in Fig. S3C, top panels). Namely, at the onset of the extinction phase (a), during extinction (b), at the onset of the renewal-test phase (c), and at the end of the renewal-test phase (d).

<u>No abrupt transition at context B onset in first sessions.</u> In the first session, the resulting cumulative response to the extinction stimulus, e.g. StimL, exhibits the expected positive slope at the end of acquisition (Fig. S3C1, black trace), which remains for several trials during the extinction phase (indicated by the gray shaded area and the horizontal red bar in Fig. S3C). This smooth transition at the onset of context B (Fig. S3C1, (a)) is driven by the strong association between StimL and the left response that was established during the acquisition phase, which is reflected in the strong net activation of the left motor unit due to the presence of StimL (Fig. S3C1 (a) in the bottom panel). As the extinction phase progresses, the conditioned choice is gradually suppressed, since the lack of reinforcements to operant responses to StimL leads to the emergence of negative associations between StimL and the left response, and between StimL and context (Fig. S3C1 (b) black and red lines in bottom panel, respectively). Since it takes several trials without reinforcements to build up the inhibition required to suppress the activation of the L response in the presence of StimL, our model predicts that the choice behavior changes smoothly at the onset of the extinction phase in the first session.

Preference for the alternative choice during extinction. As extinction learning progresses, the model favors the alternative choice over omissions (Fig. S3C1, (b)), just as pigeons often did (see Fig. 1B, 2A). This behavior results from the higher activation of motor unit R in the presence of the extinction stimulus StimL, which is explained as follows: In our experimental paradigm, during the extinction phase, responses of the motor unit R have been reinforced in the presence of both StimR (non-extinction stimulus) and the extinction context (red illumination). Consequently, a positive association between context and the right motor unit (Fig. S3C1 (b), red box) has formed. This association between the context and the motor unit R alone is now strong enough to tilt the balance between the two responses in favor of the alternative (right) choice in the presence of the extinction stimulus StimL. As the extinction phase progresses, responses in the presence of StimL remain unrewarded. Therefore, all the inhibitory connections from StimL and context to both, the left and right motor units, are further reinforced. This example illustrates the principle behind how alternative choices arise in our model. First, there is competition between excitatory and inhibitory drive to execute a particular response. Second, the different response options (R or L) compete against one another. The model's choice is ultimately the outcome of these two levels of competition in the decision-making process.

Abrupt changes of behavior at the onset of the extinction phase. In the second session, during acquisition, a new pair of stimuli, $StimL_2$ and $StimR_2$, are associated with the left and right responses, respectively, and $StimL_2$ was chosen as the extinction stimulus. This time, upon the onset of the extinction phase, there is a positive association between context and the motor unit R, which was established in the previous session (Fig. S3C2 (a), bottom; compare with the corresponding point in Fig. S3C1). As a result, in this example, the net activation of L and R is nearly balanced, and the given response is mostly determined by the noise. In the general case, however, it is also possible that the activation of R stemming from the positive influence of context is able to tilt the balance in favor of the alternative choice (the right response in this example). In any case, an abrupt change in behavior upon the onset of context B ensues (Fig. S3C2 (a)). Since this mechanism requires previous exposure to the extinction context, this behavior could not be observed during the first session in the model, similar to our finding in pigeons (Fig. 3C).

Intermittence of the renewal effect. In sessions 1 and 2, it is possible to see how operant responses to the extinction stimulus suddenly re-emerge upon the onset of the renewal-test phase (Fig. S3C1 and S3C2 (c)). Here, renewal emerges due to the release of inhibition by context B on a specific response (Fig. S3C1 and S3C2, bottom (c)), as previously suggested (Todd 2013; Todd et al. 2014). However, this effect vanishes in the third session (Fig. S3C3, top (c), and reappears in the fourth (Fig. S3C4, top, (c)). This intermittence of the renewal-effect is explained as follows: In the third session (Fig. S3C3), $StimR_3$ is chosen as the extinction stimulus. At the onset of extinction, there is a very strong activation of the right response (Fig. S3C3, bottom (b)) due to its positive association to both $StimR_3$ and context B, established during acquisition and previous sessions, respectively. Therefore, it takes more trials to extinguish the association between $StimR_3$ and the right response, which translates to a persistent extinction curve (Fig. S3C3, top, gray area). This long process of extinction, in turn, results in a very strong negative association between $StimR_3$ and the right response at the end of the extinction phase (Fig. S3C3, bottom (b)). Therefore, at the onset of the renewal-test, the release of context inhibition on the right response is not sufficient to drive the renewal effect (Fig. S3C3, bottom (c). In the fourth session, however, the strength of both negative and positive associations between context and the right response are close to their balanced saturated state (Fig. S3C4, bottom, (a)). Thus, the extinction process increases only the negative association between $StimR_4$ and the right response, leaving the remaining context-right response associations intact (Fig. S3C4, bottom, (b)). In this case, at the onset of the renewal-test phase, the net input to the right response resulting from the negative and positive associations between $StimR_4$ and the right response established during acquisition and extinction is still positive, resulting in the re-emergence of renewal (Fig. S3C4, (c)).

In the example sequence shown in Figure S3C, the associations between the extinction context and the motor response that is no longer rewarded during extinction created imbalances at the initial stage of extinction (Fig. S3C, points a). These imbalances gave rise to counterintuitive behaviors in sessions 1 to 3. However, once both positive and negative associations between the context and L and R responses balance each other out, the context can no longer exert its counter-intuitive effect on the responses. Such wearing-out of the context effectiveness not only predicts a more prominent preference for alternative choices during the first session, but also an overall decay of the renewal effect.

**Different rates of extinction learning.** Extinction in our model occurs at different speeds in different sessions (Fig. S3), even though the learning rate parameter in our model is held fixed across sessions. The varying extinction speed is an emerging phenomenon that is driven by two factors. First, the learning rule adjusts weights proportionally to the difference between the current weight and the saturation weight. Since weights are maintained across sessions, they start at different levels in different sessions. As a result, the weight adjustments, and in turn the learning speeds, differ. Second, the emitted response is determined in a competition between the influences of the stimulus and that of the context. Combined with the first point, the associative strength depends on the history of previous and concurrent exposures. For instance, if the context had not previously acquired much associative strength, the conditioned response can be suppressed quite quickly by adding negative associative strength to the context (Fig. S3 C1). If on the other hand, the context's overall associative strength was positive and its magnitude higher than that of the overall negative associative strength of the extinction stimulus initially, the model continues to emit conditioned response for a longer period during extinction until they are suppressed by negative associations with the context (Fig. S3 C3).
